# Whole-brain normative modeling across multiple functional connectivity metrics in MEG

**DOI:** 10.64898/2026.10.21.760411

**Authors:** Janus R. L. Kobbersmed, Chetan Gohil, Mark Woolrich, Diego Vidaurre

**Affiliations:** Department of Clinical Medicine, Center of Functionally Integrative Neuroscience, Aarhus University, Universitetsbyen 3, 8000 Aarhus, Denmark; Research Department of Epilepsy, UCL Queen Square Institute of Neurology, University College London, United Kingdom; Oxford Centre for Human Brain Activity, Oxford Centre for Integrative Neuroimaging, University of Oxford, Oxford, United Kingdom; Centre de Recerca Matemàtica, Campus UAB, Carrer de l’Albareda, Edifici C, 08193 Bellaterra, Barcelona, Spain

**Keywords:** MEG functional connectivity, normative modeling, network models

## Abstract

Normative modeling has recently emerged in structural and functional neuroimaging as a promising approach to characterize neuropathology at the individual level. Trained on large healthy cohorts, these models predict expected measurements from covariates such as age and sex while quantifying normal population variability. Individual measurements can then be compared against the model to identify deviations associated with pathology. While most normative models of brain function have been based on functional MRI, EEG and MEG offer more direct measurements of brain activity at higher temporal resolution. Despite the promise, how to adequately perform normative modelling at the whole-brain level on these modalities remains challenging. We here propose M/EEG-FUNCOIN to identify age- and sex-related dimensions of change in functional connectivity from a plethora of FC measures from electrophysiology: amplitude envelope correlation, phase-locking value, and time-delay embedding covariance. Using resting-state MEG data from the Cambridge Centre for Aging and Neuroscience (Cam-CAN) dataset, we show that M/EEG-FUNCOIN can: (i) characterize age- and sex-related whole-brain functional connectivity patterns in greater detail than existing methods; and (ii) effectively capture meaningful deviations in connectivity in individuals that rank high and low in cognitive ability. Altogether, these results show great potential for applications in complex, network-level brain pathology.

## 1. Introduction

Brain function biomarkers remain scarce for many brain diseases and disorders. Identifying objective indicators of dysfunction is challenging because these conditions often lie on a continuous spectrum, patient groups are heterogeneous, and large cohorts are often difficult to recruit (Calhoun et al., 2021; Moriarity et al., 2022). Recently, normative models have gained popularity in neuroimaging because they address these challenges by utilizing large cohorts of healthy subjects to characterize the healthy variation of measurements given demographic covariates such as sex and age (Fraza et al., 2025). Modeling the association to such known covariates removes variance attributable to demographic differences, making the remaining variance reflect other aspects of individual differences such as natural, healthy variation and, potentially, pathology. Based on this residual variance, potential pathology is indicated by subject-level deviations from the norm (reference curve) that are greater than would be expected from chance alone.

While most applications have focused on structural and functional MRI (s- and fMRI) (Marquand et al., 2019), normative modeling of magneto- and electroencephalography (MEG and EEG) data remains limited. In contrast to fMRI, M/EEG measures brain activity directly, has high temporal resolution, and allows for frequency-resolved characterization of brain activity (Brookes et al., 2011). Crucial for the development of normative models that embrace these qualities, large, high-quality M/EEG datasets from healthy subjects have recently been released. However, compared to s- and fMRI, M/EEG poses additional modeling challenges because of the complexity of the signals —e.g. the presence of nested oscillations at a broad range of frequencies. This motivates a plethora of measures of functional connectivity (FC) beyond covariance or correlation —which are standard in fMRI— to model the various forms of coupling between sensors or sources.

In M/EEG, the most common measures subject to normative modeling are non-FC measures such as power spectral density, magnitude, or peak frequency within bands (Italinna et al., 2023; Janiukstyte et al., 2023; Lefebvre et al., 2018; Owen et al., 2023; Zamanzadeh et al., 2026). The few previous normative models of electrophysiological FC are based on EEG data and include modeling of phase coupling (Dong et al., 2025) and amplitude envelope correlation (AEC)(Ebadi et al., 2025; Tabbal et al., 2025). One of these studies used normative modeling of average AEC across connections (Tabbal et al., 2025), while others have used edge-wise models, i.e. modeling the individual FC connections separately (Dong et al., 2025; Ebadi et al., 2025; Tabbal et al., 2025). These previous edge-wise or averaged normative models of FC in EEG identified limited effects of age on the functional connectome either after averaging or when correcting for the large number of connection models proposed (Dong et al., 2025; Ebadi et al., 2025; Tabbal et al., 2025). We suggest that models that capture a stronger association with covariates may perform better in clinical or research applications, because the residual variance reflects, to a larger degree, relevant aspects of inter-subject variation. Furthermore, edge-wise approaches overlook systemic effects (Kobbersmed et al., 2026), despite growing evidence that many higher-order functions as well as disease phenotypes arise not only from properties of focal connections but also from functional networks and, in the case of diseases, network-level consequences of localized pathology (Zhang et al., 2021).

To address these challenges, we investigate how to effectively perform whole-brain normative modeling of M/EEG FC, capturing network-level aging effects on MEG FC. Specifically, we extend the FUNCOIN framework —which we recently proposed as a tool for normative modelling of whole-brain FC in fMRI (Kobbersmed et al., 2026)—to electrophysiology. M/EEG-FUNCOIN is a normative modeling framework for M/EEG FC based on performing simultaneous dimensionality reduction and regression. This allows for modeling FC with a set of relevant components, thereby moving the modeling to the whole-brain level and avoiding the mass hypothesis testing generated by connection-wise, localized models. Our approach includes a broad range of FC measures capturing different aspects of connectivity: AEC (Hipp et al., 2012), phase locking value (PLV)(Fries, 2015; Lachaux et al., 1999), and time-delay embedding (TDE), which is a way to characterize both power and phase relations from broad-frequency data (Vidaurre et al., 2018).

To evaluate M/EEG-FUNCOIN, we use resting-state MEG data from healthy subjects in the publicly available Cam-CAN dataset (612 subjects, aged 18-88 years, scanned while lying with their eyes closed and not performing any particular task). The models revealed whole-brain FC aging effects that could not be found in previous connection-wise studies of resting-state FC in MEG (Coquelet et al., 2017) or by deriving models of average FC as previously suggested in EEG (Tabbal et al., 2025). Finally, based on a classification task on cognitive performance, we test the ability of the normative models across FC measures and frequency bands to characterize performance in a broad battery of cognitive tests.

## 2. Methods

### 2.1. Data

We use data from the Cambridge Centre for Ageing and Neuroscience (Cam-CAN) dataset, consisting of 612 healthy subjects aged 18-88 years and including resting-state MEG scans, demographic variables, structural MRI scans, and scores from a battery of cognitive tests (Shafto et al., 2014; Taylor et al., 2017). MEG recordings were obtained for 8 minutes while the subjects were awake with their eyes closed without performing any task. The initial training and testing of the M/EEG-FUNCOIN and average-FC models (Figs. 1-3) were performed on 500 training subjects (male proportion 0.508, mean age 55.41 years) and 112 held-out subjects (male proportion 0.500, mean age 53.38 years) (Table 1). The data are publicly available upon application, and no separate ethics approval was needed for this study.

**Figure 1:**
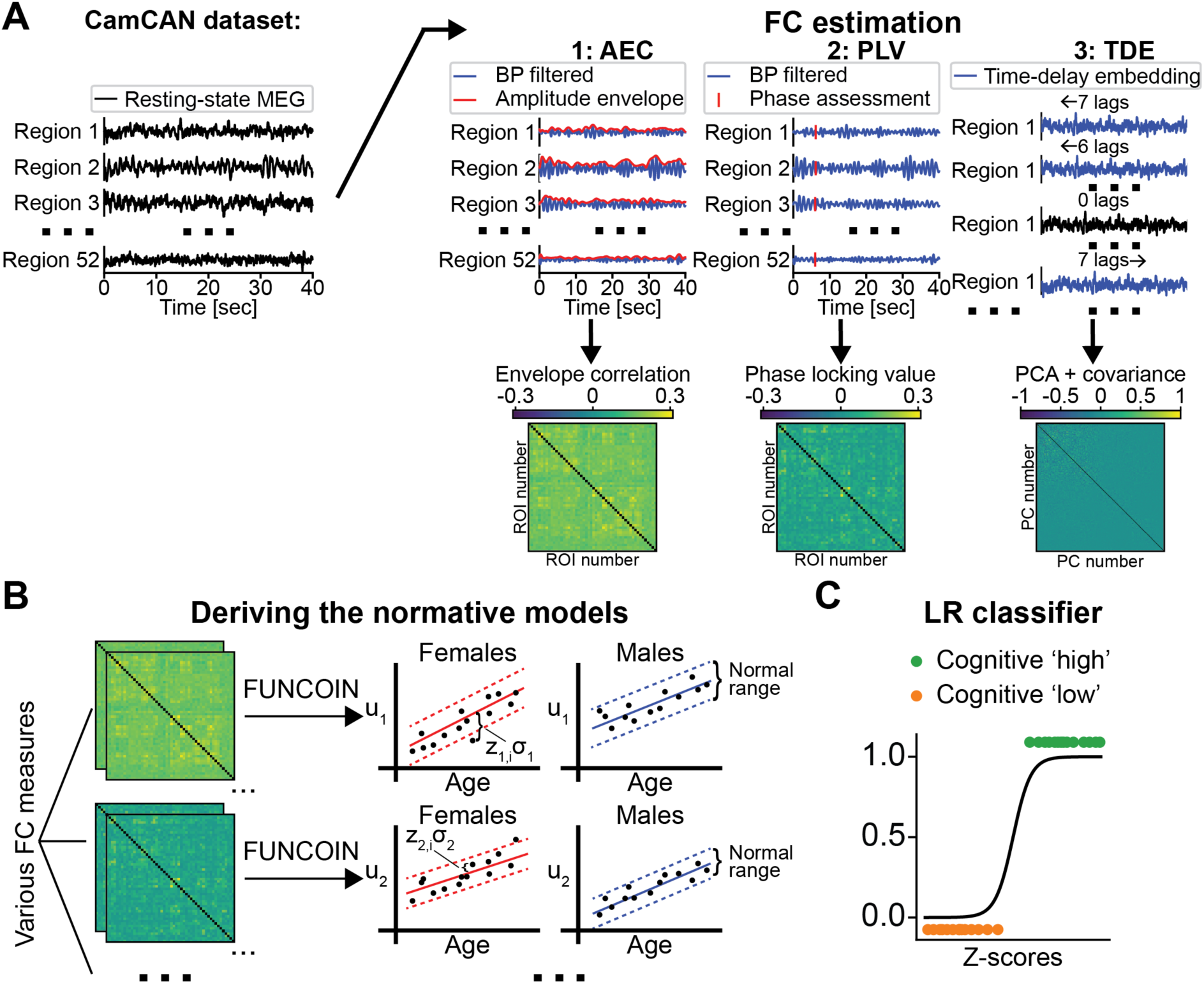
Schematic illustration of the M/EEG-FUNCOIN. **A.** Using preprocessed, source-reconstructed MEG data, we derive three FC measures: Amplitude envelope correlation (left), phase locking value (middle), and covariance after time-delay embedding (right). **B.** M/EEG-FUNCOIN identifies components from the FC matrices that are shared among the training subjects and whose strengths are maximally associated with sex and age. This forms the basis for the normative models. **C.** The ability of the Z-scores to capture higher-order brain function is assessed with a logistic regression (LR) classifier predicting cognitive test performance from the Z-scores.

**Figure 2:**
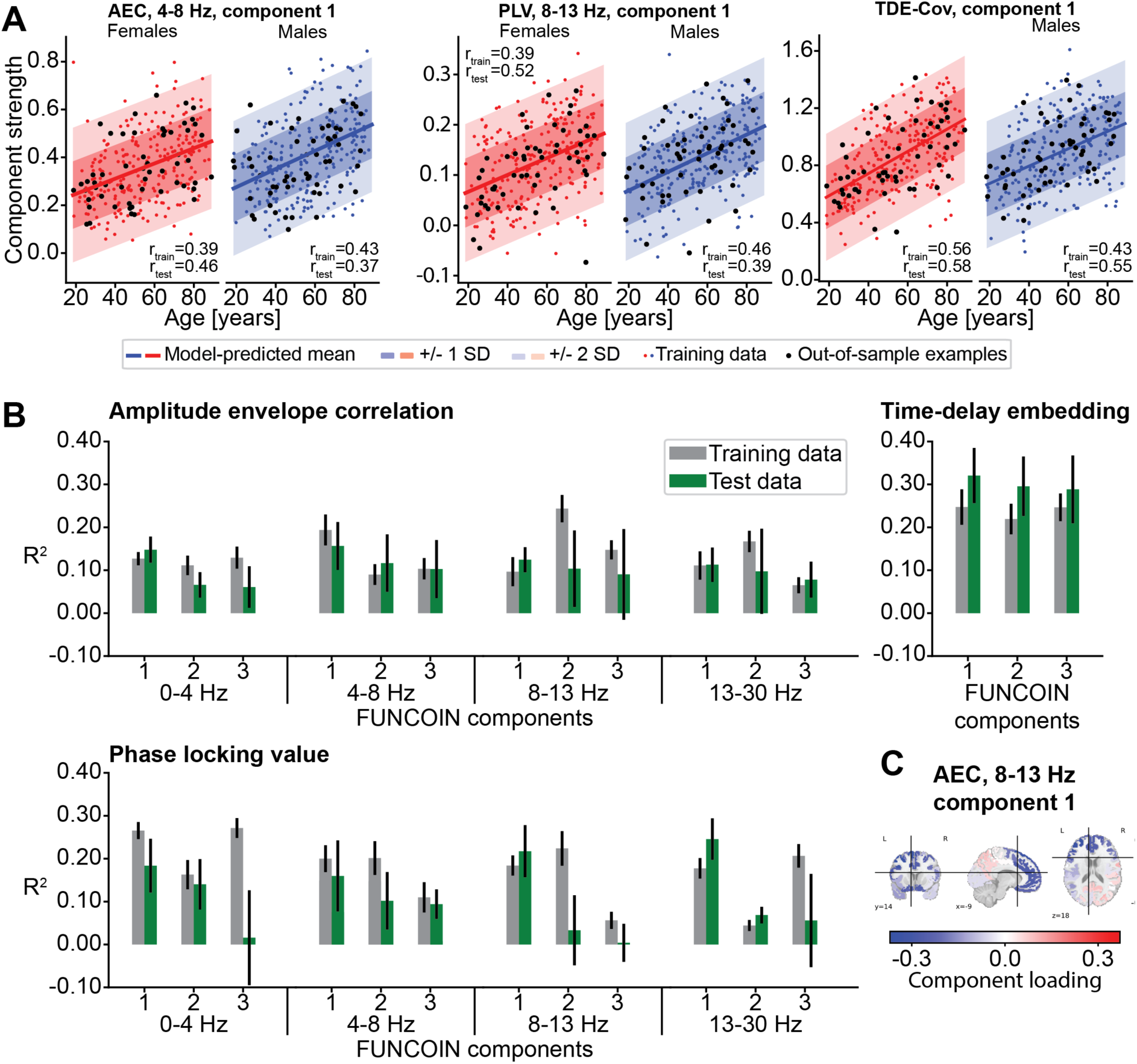
Normative models of resting-state MEG FC. **A.** Examples of normative graphs for the three different FC measures: Theta band AEC (left), alpha band PLV (middle), and TDE (right). The examples show the strong relation between the strengths of the identified components and age. Red and blue dots represent the training data, and black dots are from out-of-sample test subjects. Lines and shadings show model prediction +/- 1 or 2 SD. **B.** R^2^ train and test values across FC measures and frequency bands from the components with the top three test R^2^. M/EEG-FUNCOIN can identify FC components explaining considerable variance associated with sex and age. This makes the M/EEG-FUNCOIN components good candidate measures for normative modeling of brain function. Error bars show ±SD estimated by bootstrapping (10,000 resamples). **C.** Example visualization of the brain map of an example M/EEG-FUNCOIN component. Inspection of component loadings and the brain maps derived from them allows for identifying the brain regions involved in the sex- and age-dependent network components.

**Figure 3:**
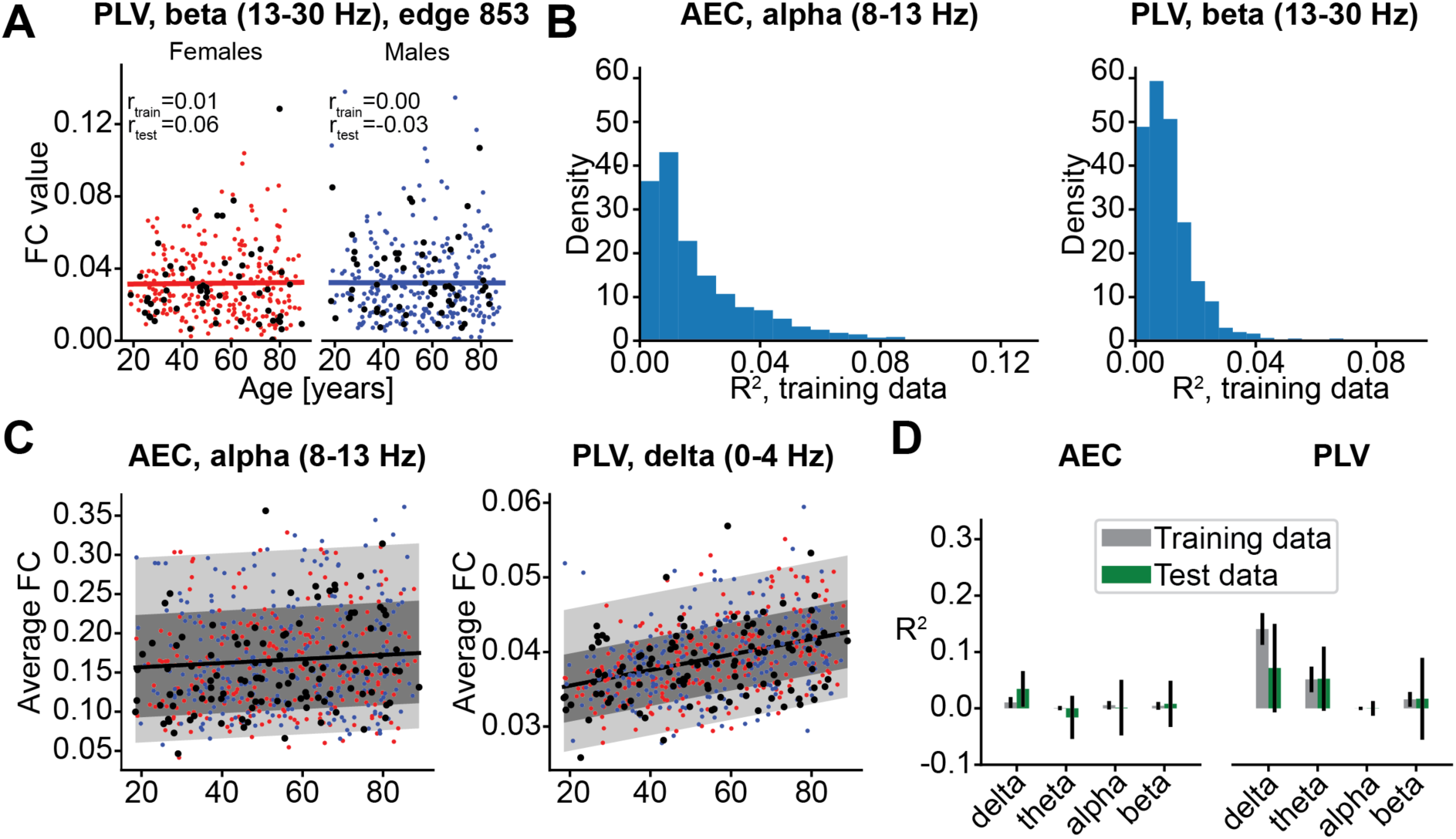
Normative models based on individual or average FC edges. **A.** Example edge from PLV in beta band, which shows no correlation between FC (PLV) values and age and sex. Red and blue dots represent the training data. Black dots are from out-of-sample test subjects. **B.** Histograms of edgewise R^2^ values after fitting the best straight line for each edge in alpha band AEC (left) and beta band PLV (right). By far the most edges show little or no change with age and sex, as witnessed by the low R^2^ values. Histograms from all frequency bands can be found in Supplementary Figure 1. **C.** Examples of normative models from averaging all FC edges in alpha band AEC (left) and delta band PLV (right). See also Supplementary Figure 2. Like in A, red and blue dots represent the training data, and black dots are from out-of-sample test subjects. **D.** R^2^ values in from train and test data show that most variance explained by age is in delta band PLV. For the other frequency bands and for AEC in general, little or no age effect was found.

**Table 1:** Demographic data.

| Subjects | N | Mean age ± SD [years] | Males (proportion) |
| --- | --- | --- | --- |
| ***CamCAN M/EEG-FUNCOIN models (Figure 2)*** |  |  |  |
| *Training* | 500 | 55.41±18.15 | 0.508 |
| *Test* | 112 | 53.38±19.02 | 0.500 |
| ***Classification task (Figure 4)*** |  |  |  |
| *Training* | 428 | 53.49±18.06 | 0.493 |
| *Test ('low')* | 92 | 58.19±19.42 | 0.674 |
| *Test ('high')* | 92 | 59.07±17.50 | 0.402 |

### 2.2. Preprocessing

The data were preprocessed as reported previously (Gohil et al., 2026). MEG data were preprocessed and source reconstructed using osl-ephys (van Es et al., 2025). Raw recordings were first MaxFiltered using temporal signal space separation (tSSS) and head motion correction, before bandpass and notch filtering and downsampling to 250 Hz. Automated detection of bad segments and channels was performed using variance-based outlier detection with the G-ESD algorithm, followed by ICA-based removal of ocular and cardiac artefacts and interpolation of bad channels. MEG data were coregistered to the structural MRI data using RHINO, and forward models were computed with a single-shell boundary element model on an 8 mm volumetric grid. Source localization was performed using a unit-noise-gain LCMV beamformer. Source-level activity was then parcellated into 52 anatomically defined ROIs (Kohl et al., 2023) by applying PCA to voxel time courses within each parcel and retaining the first principal component. To reduce spatial leakage, parcel time courses were symmetrically orthogonalized (Colclough et al., 2015), and sign ambiguities across subjects were resolved using a template-based sign-flipping procedure. Finally, parcel time courses were z-scored prior to further analysis. See the supplementary information in (Gohil et al., 2026) for full details.

### 2.3. FC estimation

We explored three different measures of functional connectivity: 1) AEC in predefined frequency bands, 2) PLV in predefined frequency bands, and 3) covariance of time-delay embedded time series.

For each subject, AEC and PLV matrices were calculated by first, for each brain region, band-pass filtering the signal and computing the analytic signal given by the Hilbert transform. For AEC, for each pair of brain regions, the correlation of the amplitudes (i.e. absolute values of the Hilbert transform) was calculated, resulting in a [no of ROIs]x[no. of ROIs] correlation matrix. For each pair of brain regions, (j,k), PLV was calculated as

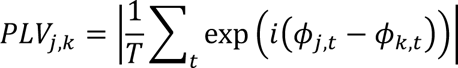

where ϕ_j,t_, ϕ_k,t_, are the phases of signals *j* and *k* at time point *t*, as obtained from the analytic signal, and T is the number of time points. The PLV takes values between 0 and 1 and quantifies the consistency of the phase difference between two brain regions over time.

TDE is based on creating time-shifted copies of the signal from each brain region in order to capture covariation despite potential phase differences. Thereby, it is a data-driven approach to characterize both amplitude and phase relationships between brain areas across frequency bands with no need for filtering (Vidaurre et al., 2018). We used a symmetric window of length 15 time points with a step size of 1 time point, which created 15 new “channels”/signals for each ROI (like in Gohil et al., 2023; Vidaurre et al., 2018). The 15 signals from the same ROI are highly correlated. We therefore performed principal components analysis by finding common principal components of the average FC matrix in the training data (N=500 in Figure 1-2), retaining the first 208 principal components (cumulative explained variance of ∼81%). The M/EEG-FUNCOIN model was then fitted to the 208×208 covariance matrices. The values of the principal components from the training data were found in the test data to calculate the TDE covariance matrices for held-out data. The same pipeline was carried out for the classification task (Figure 4), where *N_train_*=428, *N_test,low_*=*N_test,high_*=92.

M/EEG-FUNCOIN requires the FC matrices to be positive semidefinite, i.e. having non-negative eigenvalues. This is always the case for AEC and covariance of TDE. For PLV we verified this numerically for all subjects and for all frequency bands.

### 2.4. Fitting and bias removal of the FUNCOIN model

FUNCOIN and the covariance regression method, which FUNCOIN is based on (Zhao et al., 2021), were described in the first FUNCOIN paper (Kobbersmed et al., 2026). In brief, the model performs dimensionality reduction and regression in one step by finding components, ***γ*** ∈ ℝ*^p^*, of a set of *p*-by-*p* FC matrices such that the components are shared among the subjects and their strengths are most associated with covariates. This is done by optimizing the following heteroscedastic, log-linear model

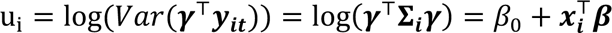

where *β_o_* ∈ ℝ is the intercept, ***β*** ∈ ℝ*^q^*^-1^ are model coefficients, and ***x_i_*** ∈ ℝ*^q^*^-1^ is the vector of *q*-1 covariates of subject *i*. The logarithm ensures that model-predicted component values respect the positive definiteness of the FC matrices.

The fitting algorithm identifies a pair of component (**γ**) and coefficients (**β**) that together minimize minus the log-likelihood function:

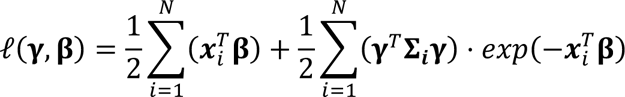

The identified component is then removed from the subjects’ FC matrices by matrix deflation before more components are identified, yielding a set of orthogonal components (Kobbersmed et al., 2026; Zhao et al., 2021). The model predictions after the log-likelihood optimization are biased (Kobbersmed et al., 2026). To remove this bias after identifying **γ** and **β**, we refit **β** by minimizing the squared residuals of the model while keeping the identified **γ** fixed.

Note that, contrary to ordinary linear regression, the best fit solution (which will be the first component identified) does not necessarily yield the highest R^2^ value. This is because the model favors components whose strength depends on the coefficients by optimizing a non-linear balance between the variance captured and the associated coefficients as seen from the loglikelihood function above. It is therefore not guaranteed that the best fit (**γ**, **β**) will yield the largest R^2^ or optimize other traditional model evaluation metrics. In practice, we therefore identify as many components as the data allow and pick the most useful for our purpose.

### 2.5. M/EEG-FUNCOIN

We recently proposed FUNCOIN as a whole-brain normative modeling framework for fMRI (Kobbersmed et al., 2026). In this paper, we studied how to adapt this method to deal with M/EEG FC metrics. To make this method suited for M/EEG, we need to address mathematical challenges related to the highly correlated nature of electrophysiological signals, which can lead to potentially rank-deficient FC matrices. We also need to ensure computational efficiency when handling large amounts of data and the high dimensionality of FC measures like TDE.

In electrophysiology, FC matrices may be rank-deficient, usually not because the data has fewer time points than signals, but because signals from, for example, sensors or time-shifted copies of signals in TDE can be highly correlated. In FUNCOIN, as proposed in fMRI (Kobbersmed et al., 2026), this would be a problem, because the fitting routine relies on matrix inversion and convergence to the optimal solution relies on identifying the minimum of a set of eigenvalues. For the inversion, we instead used the Moore-Penrose pseudoinverse. For the minimal eigenvalue, we instead used singular value decomposition and only considered the non-zero (positive) eigenvalues. This effectively restricts the analysis to the (potentially lower than full-rank) space carrying the signal. We verified numerically that these additions yielded consistent results.

High-dimensional FC matrices are frequently encountered in electrophysiology, for example in sensor space, voxel-level source-reconstructed space, or after applying TDE. The MEG/EEG-compatible version of the FUNCOIN Python package therefore contains new features for handling the additional computational load introduced by large amounts of data at a small cost of processing time. These include specifying the FC matrices with a set of eigenvectors and eigenvalues, which saves memory if the FC matrices are not full rank. Alternatively, or in combination with the eigenvector representation, the FC matrices can be “fed” to the model before fitting, which saves them in a temporary folder from where they are loaded when needed in the fitting routine. See the documentation of the FUNCOIN Python package for details on how to utilize these features.

### 2.6. Determining the number of components

When calculating FC as AEC and PLV, we first fitted the model in each frequency band to all the training data (*N*=500), identifying the 10 first component-coefficient pairs, (**γ_1_**,**β_1_**), (**γ_2_**,**β_2_**), …, (**γ_10_**,**β_10_**), while the first 6 component-coefficient pairs were found for TDE-Cov.

To test how consistently the components were identified, we then ran 10 repetitions of 10-fold cross-validation (CV) and compared the originally identified components and the 10 components identified in each of the CV folds. The order of the identified components can change between CV folds. The similarity was determined as the maximal absolute dot product between the originally identified component and the 10 components in each CV fold. Since the components are of norm 1, this is the cosine similarity. A value of 1 reflects perfect alignment, while 0 means that the components are orthogonal. For determining the number of components to keep for analysis, we chose a threshold of average similarity of 0.85 across CV folds (see Supplementary Figure 5). This yielded several M/EEG-FUNCOIN components to analyze (Supplementary Table 1), but for clarity and to limit the number of hypotheses to test, we only used the first identified M/EEG-FUNCOIN components in the classification task in Figure 4.

With TDE, the lagged signals are constructed by creating time-shifted copies of the signal from each ROI. Thus, while the M/EEG-FUNCOIN components are spatially orthogonal in the p-dimensional space of the ROIs when fitting to a p-by-p FC matrix, identifying multiple components after TDE yields orthogonal components in the *p*\*[no. of lags]-dimensional space.

### 2.7. Alternative normative models of FC

We contrasted the normative models from M/EEG-FUNCOIN to a normative model based on the average connection strength as quantified with AEC and PLV in each of the four frequency bands. Fitting a linear regression model revealed that the average FC values did not depend strongly on sex or age, except for the age dependence of average PLV in 0-4 Hz (Figure 3 and Supplementary Figure 2). In Figure 3, we therefore showed the best fit with only age as a covariate. The residuals were very skewed around the model prediction, which is not a problem for the best straight line fit, but needs to be considered when calculating the Z-scores. For the Z-scores in the cognitive classification task, we fitted a linear regression model with sex and age as covariates, followed by fitting a skew normal distribution to the residuals. Then, for each subject, we found the Z-scores based on the percentiles of the skew normal distribution. Importantly, in all frequency bands, the resulting Z-scores were standard normally distributed for both AEC and PLV, which ensures the validity of the Z-scores.

### 2.8. Classification of cognitive performance of test subjects

The Cam-CAN dataset contains neuroimaging data from cognitively healthy subjects (Mini-Mental State Examination (MMSE)>24) without neurological or serious psychiatric conditions (Taylor et al., 2017). For this reason, we could not contrast disease cases with healthy controls. Instead, we tested the ability of our model to distinguish between “high” and “low” cognitive performers, based on a wide range of cognitive tests (Shafto et al., 2014). Previous studies (Borgeest et al., 2020; Tibon et al., 2021) have used 13 cognitive scores spanning five cognitive domains (executive function, language, emotion, memory, and motor control), which were provided by Borgeest et al. Following the practice from Gohil et al (Gohil et al., 2026), three scores, being discrete or largely non-Gaussian, were excluded because they were not fit for Principal Component Analysis (PCA). This left us with 10 cognitive scores from 612 subjects (Supplementary Figure 3).

Our proof-of-concept goal was to identify ’low’ and ’high’ cognitive performers based on their Z-scores from the normative models. Since the cognitive scores decline with age (Gohil et al., 2026), we performed deconfounding on the raw cognitive scores to remove the age dependency of the cognitive scores. This was done by fitting a second-order polynomial with age as the covariate to the cognitive test scores and, for each subject, subtracting the model-predicted cognitive scores from their raw scores (see Supplementary Figure 3). Deconfounded cognitive scores were then standardized to mean 0 and variance 1 before performing PCA. The first principal component (PC) explained 29.6 % of the variance. The PC loadings revealed that better performance in any task increased the value of the first PC, and we therefore took this value as a measure of cognitive function across tasks to be used in the following classification task. Fitting a linear regression model to the resulting cognitive PC scores with age as a covariate revealed no association with age (Supplementary Figure 3, last panel).

To test whether Z-scores from M/EEG-FUNCOIN components and average FC captured differences in brain function related to cognitive function, we retrained the M/EEG-FUNCOIN and average-FC models on the middle 70 % of the subjects based on their cognitive score. The low and high 15 % were used for testing by, for different combinations of Z-scores from M/EEG-FUNCOIN and the average-FC models, training a logistic regression classifier with a small amount of regularization (C=1e5). Significance of the coefficients for each feature was assessed with permutation testing, *n_perm_*= 10,000, quantifying the proportion of the model fits to permuted data that yielded a coefficient magnitude higher than with the fit to the original (unpermuted) data.

## 3. Results

### 3.1. A whole-brain normative modeling framework for M/EEG FC

The MEG signal is temporally rich and allows for a variety of FC measures, each of which is sensitive to different aspects of the data. In a normative modeling context, this raises key questions about how we should measure resting-state M/EEG FC and how best to characterize its dependence on demographic variables like sex and age. Here, we propose M/EEG-FUNCOIN to identify whole-brain effects of sex and age for normative modelling from three frequency-aware MEG FC measures: i) AEC in predefined frequency bands, measured as the between-brain-areas correlation of amplitudes (Figure 1A, panel 1). This reflects how areas of the brain co-modulate activity in specific frequencies (Hipp et al., 2012), and is regarded one of the most reliable MEG FC measures (Colclough et al., 2016); ii) PLV in pre-defined frequency bands, which, for each pair of brain regions, quantifies the consistency of the phase difference of the signals over time (Figure 1A, panel 2), thereby measuring their frequency-specific phase synchronization (Fries, 2015; Lachaux et al., 1999); iii) time-delay embedded (TDE) covariance matrix, which characterizes power as well as phase relationships between brain areas across frequencies (Figure 1A, panel 3)(Vidaurre et al., 2018).

M/EEG-FUNCOIN (an extension of FUNCOIN originally proposed in fMRI) creates normative distributions given sex and age for each FC measure (Figure 1B). This yields the expected FC, as quantified with the FUNCOIN component strengths, for a healthy subject conditioned on sex and age, as well as normative ranges of these FC metrics across frequency bands. Having established the normative ranges of FC, we calculate Z-scores of each individual and compare the normative model to a previously suggested normative modeling approach. Finally, we test the explanatory power of the M/EEG-FUNCOIN models of the different FC measures and across frequency bands to predict cognitive performance from a number of cognitive tests, and compare it to previously suggested normative models of average FC. This is quantitatively assessed through classification of extreme performers.

### 3.2. M/EEG-FUNCOIN identifies strong normative modeling candidate measures across FC modalities and frequency bands

Fitting the M/EEG-FUNCOIN model to the three FC measures - AEC and PLV across different frequency bands as well as TDE - we identified 10 age- and sex-dependent components of AEC and PLV and 6 components in TDE. Based on how consistently the components were identified in cross-validation (CV) folds, we kept between 3 and 10 components per measure and, in the case of AEC and PLV, per frequency band (delta and 1/f, theta, alpha, and beta, see model validation below, Methods, and Supplementary Table 1).

Figure 2A shows examples of normative graphs of the first component in AEC in theta band (left), PLV in alpha band (center), and TDE (right). Across all FC measures and frequency bands, a strong correlation was found between FC and age (r≥0.37 in train and test in all plotted examples), which highlights the ability of M/EEG-FUNCOIN to distill network-level components of all three FC measures with strong associations to sex and, in particular, to age.

In normative modeling, inference is drawn from the subject-level deviations from the model predictions after relating the modeled measure to variables like sex and age. Therefore, we want to explain as much variance related to the covariates as possible. To evaluate this aspect of the M/EEG-FUNCOIN model, we quantified the explained variance from training data and held-out test data for all retained components (top three components in each setting are shown in Figure 2B). The highest single-component R^2^ values were found for TDE. Components with R^2^ values above 0.1 were found for AEC and PLV in all frequency bands. This illustrates that M/EEG-FUNCOIN (see Methods) enables us to find several sex- and age-dependent components across different FC measures and frequencies.

When applying normative models to clinical cohorts, individual subjects can be identified as outliers, possibly reflecting pathology. M/EEG-FUNCOIN allows for qualitative assessment of the functional nature of such deviations through visualization and functional interpretation of the components (example shown in Figure 2C). Thereby, in a clinical setting, specific dimensions of deviation can be linked to the clinical condition studied, which is crucial for the characterization and subtyping of heterogeneous diseases — a major goal in normative modeling. Note that the colors do not correspond to activation directly. Instead, like in PCA, increased correlation between areas of the same color/sign will increase the component value. Conversely, higher correlation between opposite colored areas decreases the component strength and vice versa.

### 3.3. Localized or averaged normative models of FC have a weak association to age and sex

Previous normative models of FC in electrophysiology have all been based on AEC or phase locking between pairs of brain regions (Dong et al., 2025; Ebadi et al., 2025; Tabbal et al., 2025). In these studies, models are fitted to each connection separately, followed by summarizing the resulting Z-scores, or by averaging all the FC values to summarize the FC to a single number before calculating the Z-scores.

Figure 3A shows the train and test FC values in an example edge from beta band PLV. The best-fitted straight lines with sex and age as covariates show that no age or sex effects are observed in this edge. Quantifying the explained variance in the 1,326 unique connection-wise models for each FC measure and frequency band shows that in both AEC and PLV most edges show very little dependence on sex and age across all frequency bands (Figure 3B, Supplementary Figure 1). For this reason, any normative framework based on modeling connections one-by-one will be based on models that do not capture a strong relation between the demographic covariates and FC.

Following Tabbal et al. (Tabbal et al., 2025), we summarized all AEC values through averaging, which we also did for PLV. We then found the best fit line while taking the skew residuals into account (Figure 3C, see Methods). For AEC, very little or no age effect was seen on averaged FC values in any frequency band, in line with the findings by Tabbal et al. (Figure 3D, left)(Tabbal et al., 2025). The same was observed for PLV, except in delta band (Figure 3C-D, right panels). Comparison of the average-FC models to the components identified with M/EEG-FUNCOIN (Figure 2) reveals that M/EEG-FUNCOIN identifies network-level age effects above and beyond the average FC strength. Thus, M/EEG-FUNCOIN identifies FC structure and development beyond what is captured with edgewise or average FC models.

### 3.4. M/EEG-FUNCOIN component Z-scores are significant predictors of cognitive performance

In this paper, we argue that whole-brain normative modeling, as implemented in M/EEG-FUNCOIN, can capture whole-brain aspects of brain function that are necessary for characterizing complex pathology. But pathology is not the only reason an individual may deviate from the norm. Since Cam-CAN does not include subjects with neurological or serious psychiatric disease (Taylor et al., 2017), here we instead asked whether the M/EEG-FUNCOIN Z-scores were related to “extreme” cases of cognitive ability (either positive or negative) and contrasted them with the average-FC models. Cam-CAN includes a set of cognitive tests spanning several cognitive domains (Supplementary Figure 3)(Shafto et al., 2014). For simplicity, we first summarized these tests in a single cognitive score, taking the age-dependent cognitive decline into account (see Methods and Supplementary Figure 3). Then, to test whether M/EEG-FUNCOIN captures information about cognitive function beyond what is captured in the average FC models, we trained the M/EEG-FUNCOIN and average FC normative models on the subjects with the middle 70 % cognitive scores, while setting aside the top and bottom 15 % for testing (Figure 4A, left). Based on these models, we calculated out-of-sample Z-scores from the low and high cognitive groups (Figure 4A, middle), and then used them as features in a logistic regression classifier (Figure 4A, right). This way, we can assess the contribution of each feature to the classification of “extreme” cases (low and high cognitive performers) having controlled for the other features. Feature significance was evaluated using FDR-corrected p-values from permutation testing (N_perm_=10,000), where p-values reflected the probability of observing a coefficient of equal or greater magnitude than in the original model.

**Figure 4:**
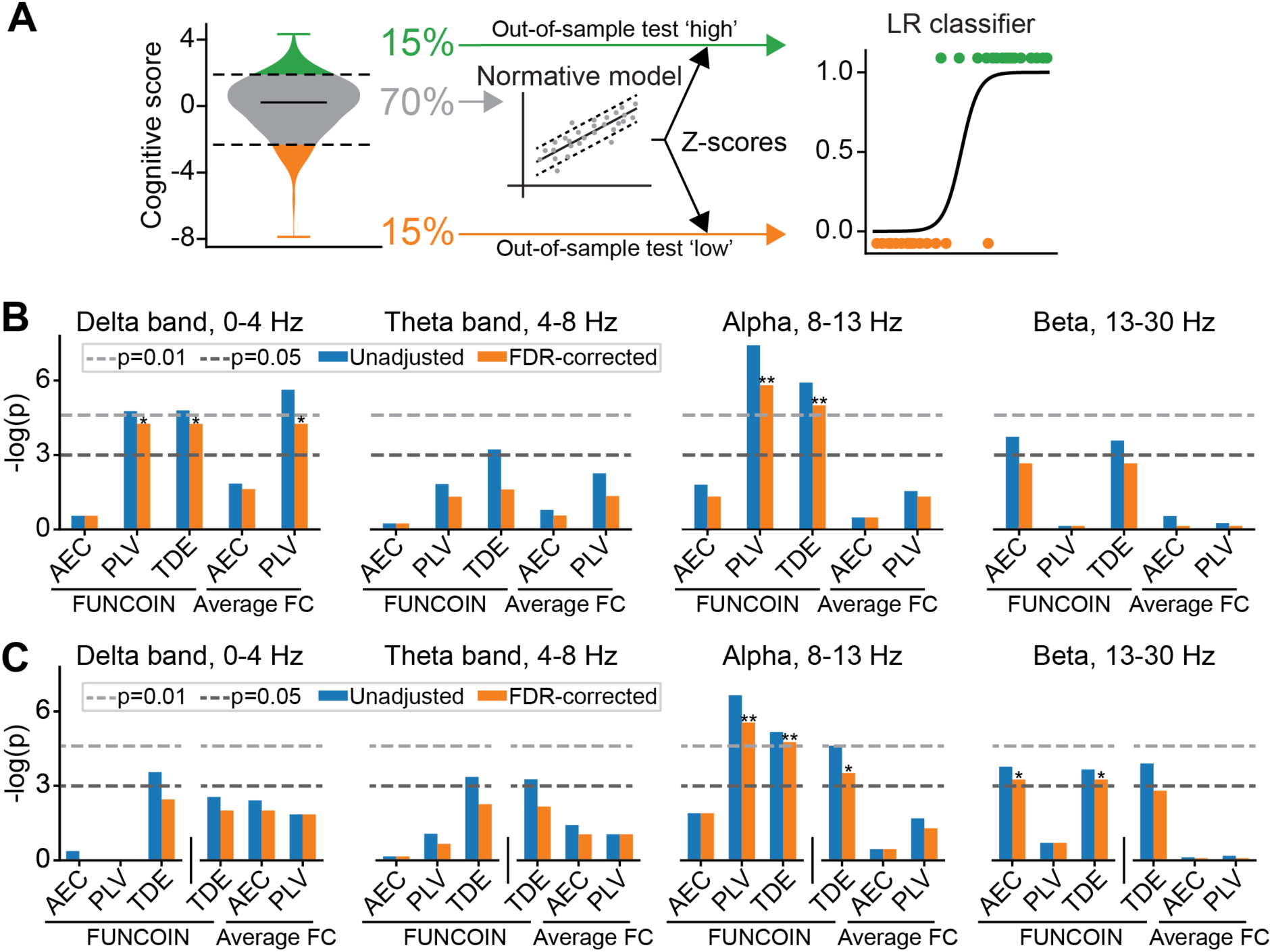
Classification of cognitive performance based on M/EEG-FUNCOIN and average-FC Z-scores. **A.** Diagram of the model pipeline for the prediction. Based on cognitive scores, subjects are split into ’controls’ (middle 70 %) and ’cases’ (bottom and top 15 %). The M/EEG-FUNCOIN and average-FC normative models are fitted on the controls. Out-of-sample Z-scores of the ’low’ and ’high’ groups are used as features in logistic regression classification to determine which Z-scores are useful for discriminating low and high cognitive performers. **B.** Frequency band-wise fitting of the logistic regression classifier and inspection of coefficient p-values reveal several candidate FC measures for normative modeling, primarily among the M/EEG-FUNCOIN components. TDE is a consistently well-performing predictor in different model settings. **C.** Coefficient p-values from fitting the classifier based on M/EEG-FUNCOIN values alone or the combination of M/EEG-FUNCOIN-TDE and average-FC. Several candidate measures from M/EEG-FUNCOIN are maintained in these restricted model settings.

First, for each frequency band, we combined the Z-scores from the first identified M/EEG-FUNCOIN component in AEC, PLV, and TDE, as well as the average FC in AEC and PLV. This yielded several candidate measures as seen from the unadjusted p-values (Figure 4B, note that -log(p) is plotted, making higher bars correspond to lower p-values). After correction for multiple hypotheses, M/EEG-FUNCOIN components from delta and alpha band PLV were significant contributors, while the TDE component had unadjusted *p*<0.05 in all settings, surviving FDR-correction in the delta and alpha band models. Average delta band PLV also made a significant contribution to the prediction.

When combining many features, the fitted coefficients and associated p-values are conditional on the rest of the included features. Therefore, a feature can become significant or not when changing the combination of features. This can be the case especially (but not exclusively) if the features are highly correlated, which was the case for the Z-scores of average- and FUNCOIN-PLV in several frequency bands. For this reason, and to reduce the number of hypotheses to correct for, we refitted the model with fewer features. Since the FUNCOIN-TDE component was a good candidate FC measure in all the tested combinations in Figure 4B, and because TDE Z-scores showed little or no correlation with any other features, we refitted the model with AEC and PLV Z-scores from *either* M/EEG-FUNCOIN or average FC in combination with the Z-scores from FUNCOIN-TDE (Figure 4C). Several M/EEG-FUNCOIN components were significant predictors of cognitive performance, including AEC in beta band, which did not survive FDR-correction in the larger model (Figure 4B). Delta band FUNCOIN-PLV went from significant to highly non-significant when omitting the average FC Z-scores. The fact that the feature was significant only conditional on the inclusion of average FC Z-scores indicates the relevance of the average FC in relation to cognition, but also that the M/EEG-FUNCOIN component provides predictive information beyond what is explained by the average FC. FUNCOIN-TDE remained a strong candidate in all feature combinations.

These analyses show the relevance and flexibility of the M/EEG-FUNCOIN components and the ability to capture higher-order aspects of brain function. Specific research questions and hypotheses will determine whether other M/EEG-FUNCOIN configurations will be relevant in that particular setting.

### 3.5. M/EEG-FUNCOIN is consistent, statistically valid, and scalable

Normative modeling relies on subject-level interpretation of the estimated Z-scores and the ability to fit to large datasets, making model validation and feasibility crucial. We therefore carefully examined the consistency of the projection estimations; statistical validity of the model fit and Z-scores; and scalability of the fitting routine.

With M/EEG-FUNCOIN, we sequentially identify multiple, covariate-related components that are shared among the subjects. The more components already identified, the greater the chance that another identified component is a result of overfitting, and not shared across training and test subjects. We therefore determined the number of components to keep for analysis based on how consistently they were estimated across CV folds in 10 repetitions of 10-fold CV (see Methods and Supplementary Table 1). In all FC measures and frequency bands, at least 3 and up to 10 components were consistently identified, meaning that M/EEG-FUNCOIN identified several dimensions of sex- and age-dependent changes in FC (Supplementary figure 4A, Supplementary Table 1).

The inference from the Z-scores relies on normally distributed residuals with homogeneous variance. We visually inspected QQ- and residual-versus-fitted plots as well as Z-score histograms for all identified components (examples in Supplementary Figure 4B). These indicated that the consistently identified component values had approximately normally distributed residuals, while residual-versus-fitted plots showed homogeneous variances (Supplementary Figure 4B, left and center). As required, the histograms of component Z-scores showed good agreement with standard normal distributions (Supplementary Figure 4B, right).

To assess scalability, we repeatedly fitted the M/EEG-FUNCOIN model to identify 2 components of the TDE covariance matrix while varying the number of subjects (N) and the number of principal components retained (p). This showed great scalability in N and p, which accommodates the derivation of normative models from large healthy cohorts and in high-dimensional cases like sensor-space data or after time-delay embedding (Supplementary Figure 4C).

Together, these findings back the appropriateness of M/EEG-FUNCOIN for normative modeling of FC in electrophysiology by showing that the models are well-posed, well-fitted, and scalable, and that the resulting Z-scores are valid.

## 4. Discussion

In this paper, we present a framework for normative modeling of FC in human whole-brain electrophysiology. For this, we proposed M/EEG-FUNCOIN, an extension of our recently developed FUNCOIN framework for fMRI (Figure 1B), to model a variety of FC metrics under minimal assumptions. We showed that M/EEG-FUNCOIN, through performing simultaneous dimensionality reduction and regression, captures sex- and age-dependent network-level effects that could not be identified using previous normative models of resting-state MEG FC. This challenges previous claims based on models of separate connections or average connectivity that the AEC connectome from resting-state MEG is preserved in healthy aging (Coquelet et al., 2017; Tabbal et al., 2025). To probe the models’ ability to capture outliers, we showed, based on classifying subjects with lowest and highest cognitive test scores, that applying M/EEG-FUNCOIN to AEC and PLV in different frequency bands, as well as to TDE–based FC, offers several good candidate FC measures for normative modeling of complex brain function.

### 4.1. Candidate FC measures for normative modeling

The strongest association between FC and age was observed with the FUNCOIN-TDE component, which also contributed significantly to the cognitive classification task. This is consistent with TDE capturing both amplitude- and phase-related aspects of the signal without relying on predefined frequency bands, thereby integrating multiple dimensions of coupling within a single representation. This makes TDE a strong candidate for normative modeling of MEG FC. In fMRI, combining FC features from different model settings has been shown to improve the prediction of individual traits (Griffin et al., 2024). This may apply to combinations of M/EEG-FUNCOIN components as well, with AEC and PLV capturing more specific aspects of FC, which the model was also able to characterize. The optimal FC representation will ultimately depend on the research question and the signal-to-noise ratio of the data (which, for example, would be lower in low-density EEG). The present study shows the versatility of the M/EEG-FUNCOIN normative modeling framework to accommodate different metrics and, thus, research questions.

### 4.2. Previous normative models in electrophysiology

Several previous normative models in M/EEG have been based on frequency band-specific power or related measures that were observed to change with age (Ebadi et al., 2025; Italinna et al., 2023; Janiukstyte et al., 2023; Owen et al., 2023; Tabbal et al., 2025; Zamanzadeh et al., 2026). While these are not measures of FC, the power of the signals can influence some FC metrics, as these are affected by the signal-to-noise ratio. Therefore, an observed age-dependent change in FC may to some degree reflect aging effects on power features. Here, we find strong aging effects on PLV, which is based on quantifying instantaneous phase relationships with minimal confounding from amplitude modulations. Thus, the normative models of PLV presented here not only move normative modeling from power characteristics to FC but also target fundamentally distinct aspects of brain function and may therefore complement existing normative approaches in clinical applications.

In agreement with earlier normative models of AEC in EEG (Tabbal et al., 2025), we found that the average AEC did not change with sex and age. Outside normative modeling, modeling of individual connections in AEC, where no significant changes were observed, has led to the conclusion that the electrophysiological connectome is maintained in healthy aging (Coquelet et al., 2017). While this is true for the average FC strength or when inspecting local connections (Figure 3), M/EEG-FUNCOIN’s network-level characterization of FC reveals age-dependent changes in AEC (and PLV), indicating that M/EEG-FUNCOIN extracts FC characteristics beyond what is captured with localized or averaged models.

### 4.3. Limitations of the study

Contrary to ordinary regression methods, where the model is fit directly to the response variables, M/EEG-FUNCOIN identifies the response variable (the components) while also fitting the model coefficients. For this reason, the fact that the identified M/EEG-FUNCOIN component strengths are well-fitted with straight lines, reflecting a linear effect of age, does not rule out nonlinear age effects. This is because components associated with other variables or variable transformations may also be identifiable. However, when fitting M/EEG-FUNCOIN with age^2^ as a covariate (results not shown), the first two components identified were the same as those identified with linear age effects as presented in this paper, and had age^2^ coefficients close to 0. This shows that the strongest effects were linear, but identifying more components or exploring other age transformations may reveal non-linear FC effects of aging; this was, however, not the scope of this paper.

In this study, we only kept the first component from each FC measure across frequency bands for the cognitive classification task. We previously found that two FC components from resting-state fMRI FC allowed for identifying distinct directions of deviation in bipolar disorder (Kobbersmed et al., 2026). Depending on the research question, more components may also be relevant to keep for analysis of FC in electrophysiology, possibly restricted to specific FC measures or frequency bands.

The features used in the classification task were Z-scores derived from FC normative models trained only on age and sex. This can be thought of as an unsupervised dimensionality reduction approach driven by the covariates; it is unsupervised because it is not using information from the final outcome of interest (here, as an example, cognitive ability). Consequently, the cognitive information available to the classifier may be limited. The goal of our analysis (Figure 4) was to identify which of the normative models contributed significantly to discriminating cognitive performance. However, a normative modeling study specifically targeted at cognitive performance or a clinical condition may be optimized in terms of choice of covariates, preprocessing steps, choice of brain parcellation, choice of FC metric, etc. For example, in fMRI, normative models have been used to correlate certain task performances with symptom severity based on existing knowledge of the disease in question (Marquand et al., 2016). The flexibility of the M/EEG-FUNCOIN model makes it straightforward to include various types of variables, categorical or continuous, to also accommodate these more hypothesis-driven normative modeling studies.

## 5. Conclusions

In this study, we present M/EEG-FUNCOIN, a whole-brain normative modeling framework for FC in electrophysiology. Fitting the model to AEC and PLV in different frequency bands, as well as TDE-based covariance, from a large, healthy cohort, M/EEG-FUNCOIN estimates sex- and age-dependent FC trajectories and establishes normative ranges across FC measures and frequency bands. Crucial for normative modeling, we demonstrate that the M/EEG-FUNCOIN framework is consistent, statistically valid, and scalable; and that it captures stronger associations between FC and demographic variables than previously suggested normative models of FC in electrophysiology. Finally, we show that components identified by M/EEG-FUNCOIN across all three FC measures are associated with cognitive ability, showing the potential of the framework for characterization and normative modeling of complex brain function.

## Declaration of the use of AI

AI was not used for planning or carrying out the research. AI (or other tools) were used to check spelling and grammar.

## Data and Code Availability

All model training and testing were done with the FUNCOIN Python package, which is available for installation via PyPI or from github.com/kobbersmed/funcoin. Data came from the Cam-CAN dataset, which is publicly available upon application. Code used for the classification task (Figure 4) will be made available on GitHub.

## CRediT authorship contributions

JRLK: Writing - original draft, writing - review and editing, formal analysis, software, visualization

CG: Writing - review and editing, resources

MW: Writing - review and editing, supervision

DV: Conceptualization, writing - review and editing, methodology, supervision

## Supplementary figures

**Supplementary Figure 1:**
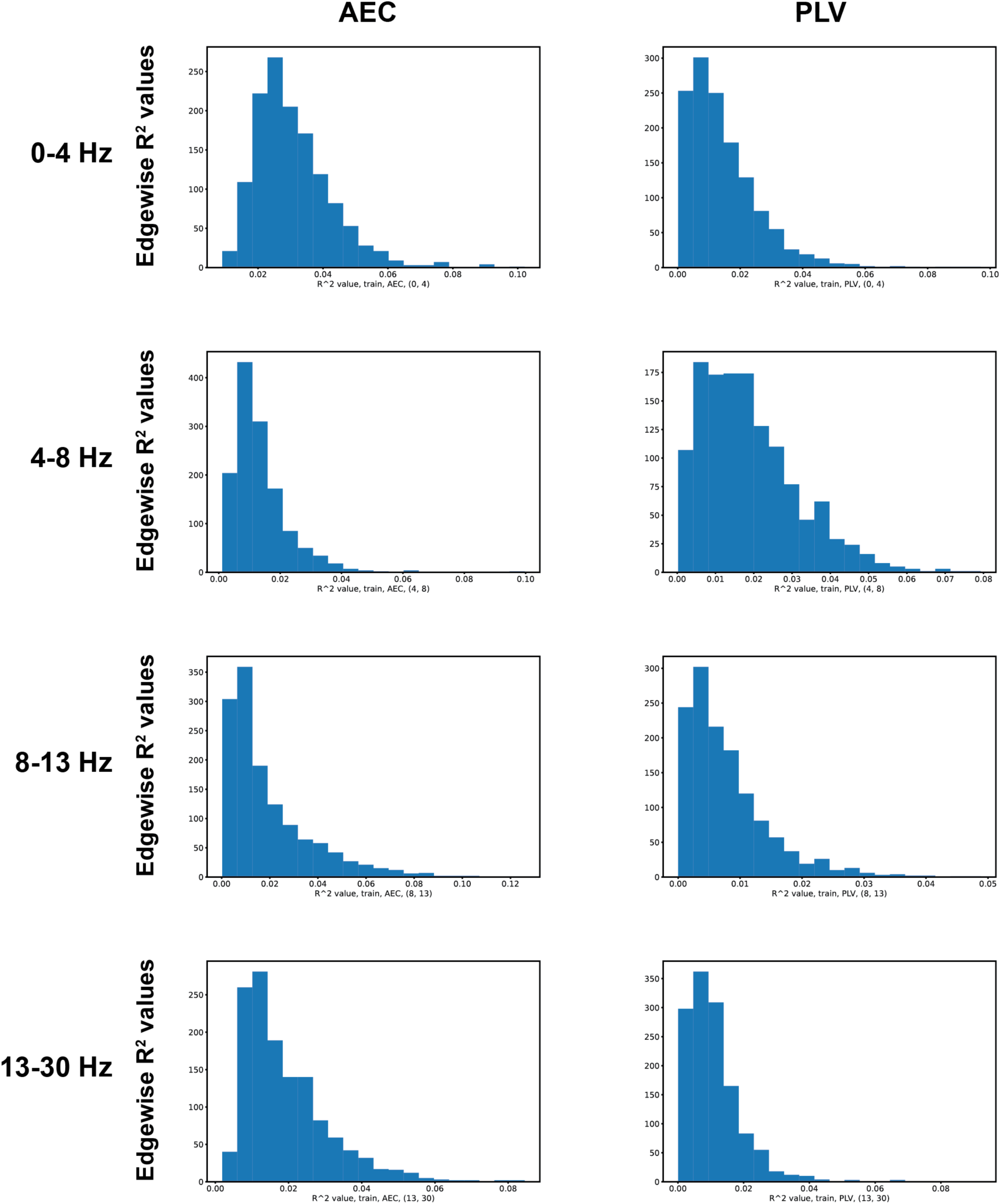
R^2^ values from edgewise linear FC models. Histograms of R^2^ after linear regression of the FC values from AEC (left column) and PLV (right column) in each frequency band (rows).

**Supplementary Figure 2:**
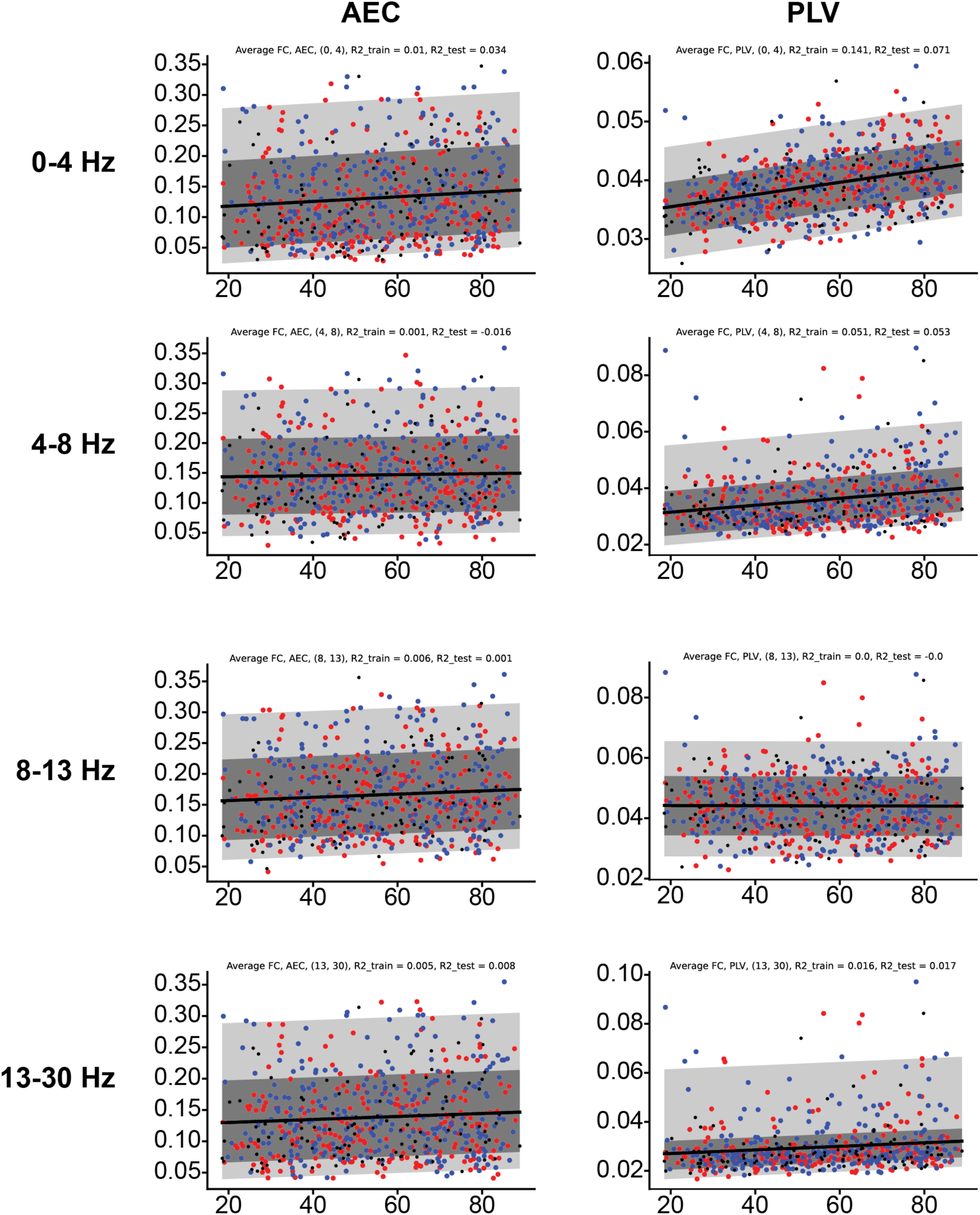
Normative models based on average FC. Normative models of average FC assessed as AEC (left column) and PLV (right column) in each frequency band (rows). The best fit straight line is shown in black. Shaded areas show percentiles corresponding to +/- Z=1 and +/- Z=2.

**Supplementary Figure 3:**
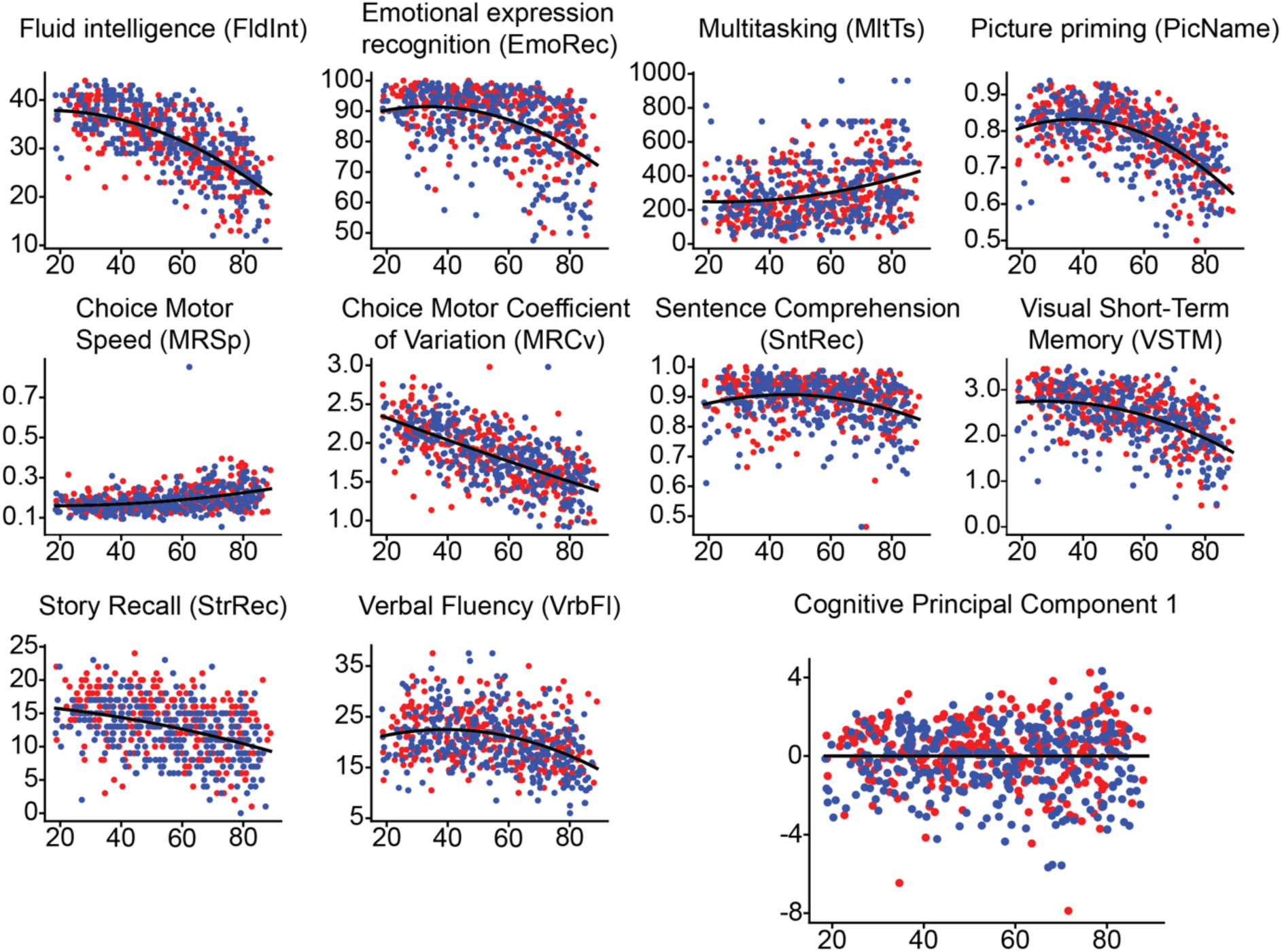
Cognitive test scroes. For each of the 10 plots included in the analysis (smaller plots), we fitted a parabola (age and age^2^ as covariates), which was used for deconfounding for age effects before conducting PCA. In the cognitive classification task (Figure 4), the first PC, which explained 29.6 % of the test score variance was used as a measure of cognitive function across tasks. No significant dependence of the PC score on age was observed.

**Supplementary Figure 4:**
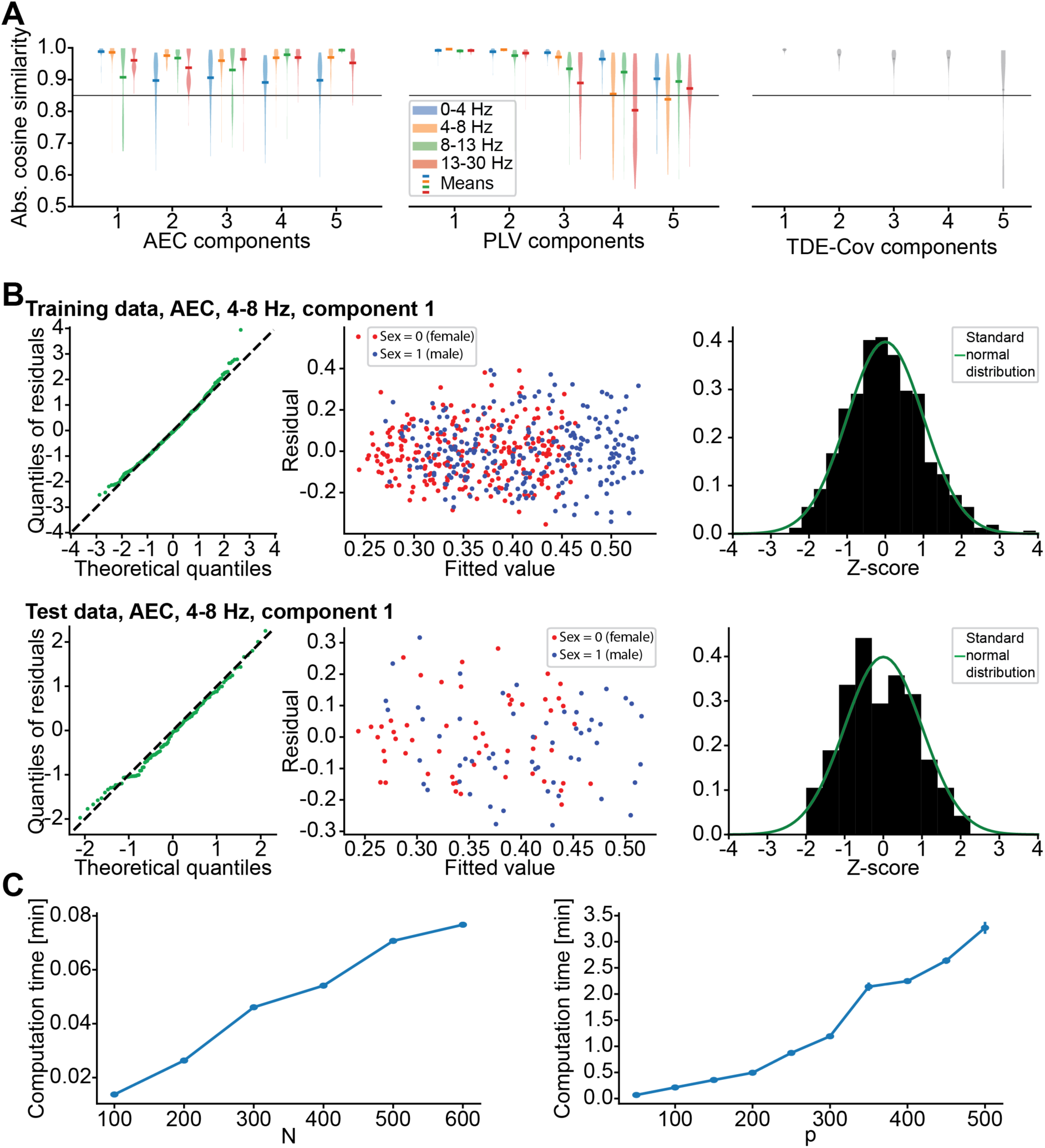
Model fit evaluation and statistical validation. **A.** Assessing consistency of the FUNCOIN component estimates revealed that several sex- and/or age-dependent components were identified in all FC measures and across frequency bands. **B.** Model validation plots in training and test data for component 1 in alpha band spectral coherence. QQ-plots (left) show normally distributed residuals; residual-versus-fitted plots (middle) show variance homogeneity; and histograms of Z-scores do not indicate deviation from a standard normal distribution. **C.** Testing scalability by repeatedly identifying 2 FUNCOIN components for varying N and p. FC was the covariance after TDE, retaining *p* principal components. Varying *N* between 100 and 600 while fitting to FC from 50 principal components after TDE showed good scalability in *N*. Varying *p*, the number of retained principal components after TDE, between 50 and 500 while fitting with *N*=500 showed good scalability in p.

**Supplementary Table 1:**
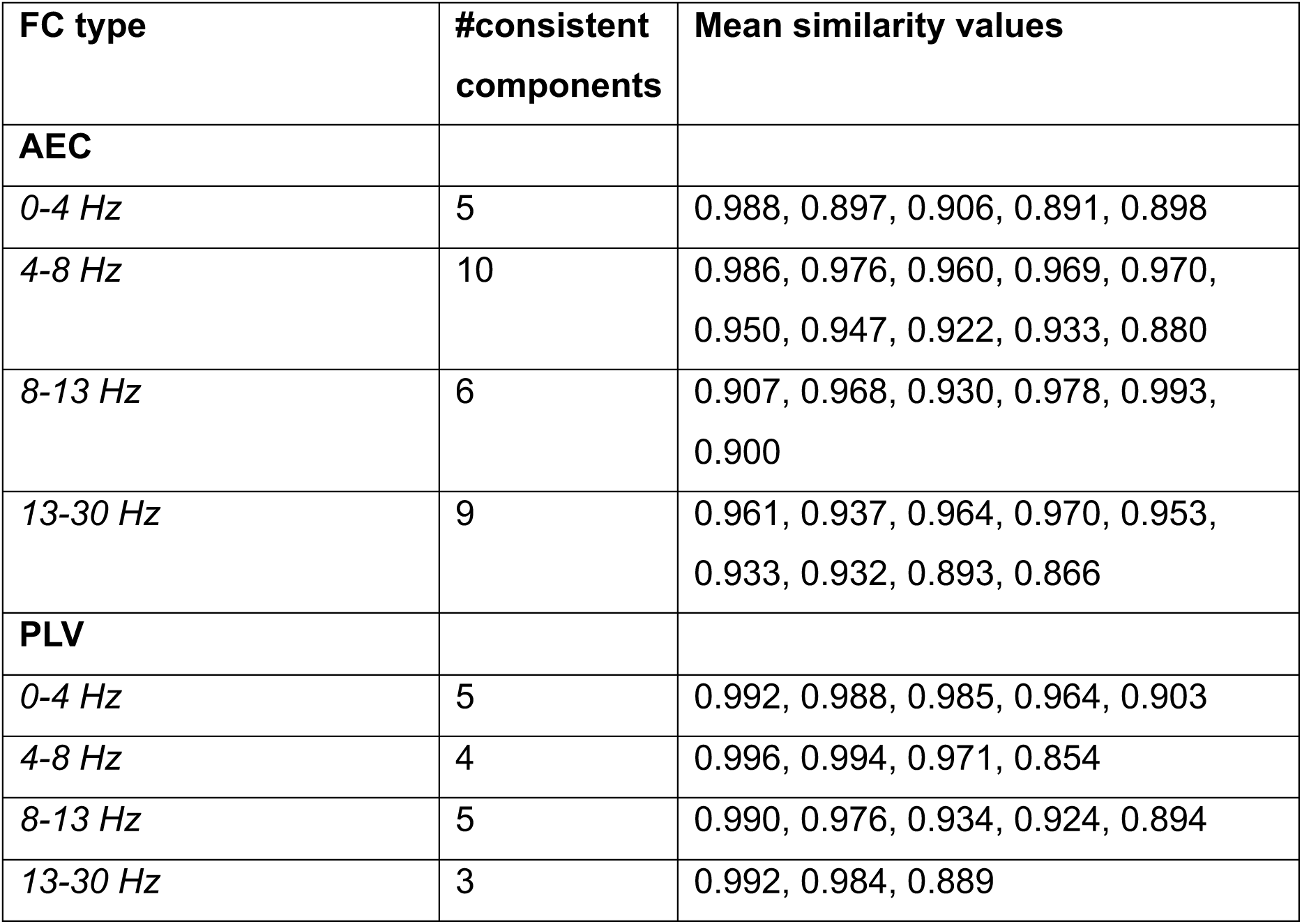
Number of FUNCOIN components consistently identified.

| **FC type** | **#consistent components** | **Mean similarity values** |
| --- | --- | --- |
| **AEC** |  |  |
| *0-4 Hz* | 5 | 0.988, 0.897, 0.906, 0.891, 0.898 |
| *4-8 Hz* | 10 | 0.986, 0.976, 0.960, 0.969, 0.970, 0.950, 0.947, 0.922, 0.933, 0.880 |
| *8-13 Hz* | 6 | 0.907, 0.968, 0.930, 0.978, 0.993, 0.900 |
| *13-30 Hz* | 9 | 0.961, 0.937, 0.964, 0.970, 0.953, 0.933, 0.932, 0.893, 0.866 |
| **PLV** |  |  |
| *0-4 Hz* | 5 | 0.992, 0.988, 0.985, 0.964, 0.903 |
| *4-8 Hz* | 4 | 0.996, 0.994, 0.971, 0.854 |
| *8-13 Hz* | 5 | 0.990, 0.976, 0.934, 0.924, 0.894 |
| *13-30 Hz* | 3 | 0.992, 0.984, 0.889 |

## References

Borgeest, G. S., Henson, R. N., Shafto, M., Samu, D., Cam, C. A. N., & Kievit, R. A. (2020). Greater lifestyle engagement is associated with better age-adjusted cognitive abilities. PLoS One, 15(5), e0230077. 10.1371/journal.pone.0230077

Brookes, M. J., Woolrich, M., Luckhoo, H., Price, D., Hale, J. R., Stephenson, M. C., Barnes, G. R., Smith, S. M., & Morris, P. G. (2011). Investigating the electrophysiological basis of resting state networks using magnetoencephalography. Proc Natl Acad Sci U S A, 108(40), 16783–16788. 10.1073/pnas.1112685108

Calhoun, V. D., Pearlson, G. D., & Sui, J. (2021). Data-driven approaches to neuroimaging biomarkers for neurological and psychiatric disorders: emerging approaches and examples. Curr Opin Neurol, 34(4), 469–479. 10.1097/WCO.0000000000000967

Colclough, G. L., Brookes, M. J., Smith, S. M., & Woolrich, M. W. (2015). A symmetric multivariate leakage correction for MEG connectomes. Neuroimage, 117, 439–448. 10.1016/j.neuroimage.2015.03.071

Colclough, G. L., Woolrich, M. W., Tewarie, P. K., Brookes, M. J., Quinn, A. J., & Smith, S. M. (2016). How reliable are MEG resting-state connectivity metrics? Neuroimage, 138, 284–293. 10.1016/j.neuroimage.2016.05.070

Coquelet, N., Mary, A., Peigneux, P., Goldman, S., Wens, V., & De Tiege, X. (2017). The electrophysiological connectome is maintained in healthy elders: a power envelope correlation MEG study. Sci Rep, 7(1), 13984. 10.1038/s41598-017-13829-8

Dong, Q., Zhou, Y., Xiong, X., Liu, P., Li, J., Luo, C., Gong, D., Dong, L., & Yao, D. (2025). A normative model-based assessment framework for large-scale, multi-site EEG data. Brain Res Bull, 231, 111546. 10.1016/j.brainresbull.2025.111546

Ebadi, A., Allouch, S., Mheich, A., Tabbal, J., Kabbara, A., Robert, G., Lefebvre, A., Iftimovici, A., Rodriguez-Herreros, B., Chabane, N., & Hassan, M. (2025). Beyond homogeneity: charting the landscape of heterogeneity in neurodevelopmental and psychiatric electroencephalography. Transl Psychiatry, 15(1), 223. 10.1038/s41398-025-03441-0

Fraza, C., Rutherford, S., Buckova, B. R., Beckmann, C. F., & Marquand, A. F. (2025). The promise of quantifying individual risk for brain disorders through normative modeling, a narrative review. Neurosci Biobehav Rev, 176, 106284. 10.1016/j.neubiorev.2025.106284

Fries, P. (2015). Rhythms for Cognition: Communication through Coherence. Neuron, 88(1), 220–235. 10.1016/j.neuron.2015.09.034

Gohil, C., Huang, R., Roberts, E., van Es, M. W. J., Quinn, A. J., Vidaurre, D., & Woolrich, M. W. (2023). osl-dynamics: A toolbox for modelling fast dynamic brain activity. In: eLife Sciences Publications, Ltd.

Gohil, C., Kohl, O., Pitt, J., van Es, M. W. J., Quinn, A. J., Vidaurre, D., Turner, M. R., Nobre, A. C., & Woolrichi, M. W. (2026). Effects of Age on Resting-State Cortical Networks. Hum Brain Mapp, 47(5), e70516. 10.1002/hbm.70516

Griffin, B., Ahrends, C., Gohil, C., Alfaro-Almagro, F., Woolrich, M. W., Smith, S. M., & Vidaurre, D. (2024). Stacking models of brain dynamics to improve prediction of subject traits in fMRI. Imaging Neuroscience, 2, 1–22. 10.1162/imag_a_00267

Hipp, J. F., Hawellek, D. J., Corbetta, M., Siegel, M., & Engel, A. K. (2012). Large-scale cortical correlation structure of spontaneous oscillatory activity. Nat Neurosci, 15(6), 884–890. 10.1038/nn.3101

Italinna, V., Kaltiainen, H., Forss, N., Liljestrom, M., & Parkkonen, L. (2023). Using normative modeling and machine learning for detecting mild traumatic brain injury from magnetoencephalography data. PLoS Comput Biol, 19(11), e1011613. 10.1371/journal.pcbi.1011613

Janiukstyte, V., Owen, T. W., Chaudhary, U. J., Diehl, B., Lemieux, L., Duncan, J. S., de Tisi, J., Wang, Y., & Taylor, P. N. (2023). Normative brain mapping using scalp EEG and potential clinical application. Sci Rep, 13(1), 13442. 10.1038/s41598-023-39700-7

Kobbersmed, J. R. L., Gohil, C., Marquand, A., & Vidaurre, D. (2026). Normative modeling of brain function abnormalities in complex pathology requires a whole-brain approach. Prog Neurobiol, 260, 102906. 10.1016/j.pneurobio.2026.102906

Kohl, O., Woolrich, M., Nobre, A., & Quinn, A. (2023). Glasser52: a parcellation for MEG-analysis. Zenodo. 10, 5281.

Lachaux, J. P., Rodriguez, E., Martinerie, J., & Varela, F. J. (1999). Measuring phase synchrony in brain signals. Hum Brain Mapp, 8(4), 194–208. 10.1002/(sici)1097-0193(1999)8:4<194::aid-hbm4>3.0.co;2-c

Lefebvre, A., Delorme, R., Delanoe, C., Amsellem, F., Beggiato, A., Germanaud, D., Bourgeron, T., Toro, R., & Dumas, G. (2018). Alpha Waves as a Neuromarker of Autism Spectrum Disorder: The Challenge of Reproducibility and Heterogeneity. Front Neurosci, 12, 662. 10.3389/fnins.2018.00662

Marquand, A. F., Kia, S. M., Zabihi, M., Wolfers, T., Buitelaar, J. K., & Beckmann, C. F. (2019). Conceptualizing mental disorders as deviations from normative functioning. Mol Psychiatry, 24(10), 1415–1424. 10.1038/s41380-019-0441-1

Marquand, A. F., Rezek, I., Buitelaar, J., & Beckmann, C. F. (2016). Understanding Heterogeneity in Clinical Cohorts Using Normative Models: Beyond Case-Control Studies. Biol Psychiatry, 80(7), 552–561. 10.1016/j.biopsych.2015.12.023

Moriarity, D. P., Joyner, K. J., Slavich, G. M., & Alloy, L. B. (2022). Unconsidered issues of measurement noninvariance in biological psychiatry: A focus on biological phenotypes of psychopathology. Mol Psychiatry, 27(3), 1281–1285. 10.1038/s41380-021-01414-5

Owen, T. W., Schroeder, G. M., Janiukstyte, V., Hall, G. R., McEvoy, A., Miserocchi, A., de Tisi, J., Duncan, J. S., Rugg-Gunn, F., Wang, Y., & Taylor, P. N. (2023). MEG abnormalities and mechanisms of surgical failure in neocortical epilepsy. Epilepsia, 64(3), 692–704. 10.1111/epi.17503

Shafto, M. A., Tyler, L. K., Dixon, M., Taylor, J. R., Rowe, J. B., Cusack, R., Calder, A. J., Marslen-Wilson, W. D., Duncan, J., Dalgleish, T., Henson, R. N., Brayne, C., & Matthews, F. E. (2014). The Cambridge Centre for Ageing and Neuroscience (Cam-CAN) study protocol: a cross-sectional, lifespan, multidisciplinary examination of healthy cognitive ageing. BMC Neurol, 14, 204. 10.1186/s12883-014-0204-1

Tabbal, J., Ebadi, A., Mheich, A., Kabbara, A., Guntekin, B., Yener, G., Paban, V., Gschwandtner, U., Fuhr, P., Verin, M., Babiloni, C., Allouch, S., & Hassan, M. (2025). Characterizing the heterogeneity of neurodegenerative diseases through EEG normative modeling. NPJ Parkinsons Dis, 11(1), 117. 10.1038/s41531-025-00957-6

Taylor, J. R., Williams, N., Cusack, R., Auer, T., Shafto, M. A., Dixon, M., Tyler, L. K., Cam, C., & Henson, R. N. (2017). The Cambridge Centre for Ageing and Neuroscience (Cam-CAN) data repository: Structural and functional MRI, MEG, and cognitive data from a cross-sectional adult lifespan sample. Neuroimage, 144(Pt B), 262–269. 10.1016/j.neuroimage.2015.09.018

Tibon, R., Tsvetanov, K. A., Price, D., Nesbitt, D., Can, C., & Henson, R. (2021). Transient neural network dynamics in cognitive ageing. Neurobiol Aging, 105, 217–228. 10.1016/j.neurobiolaging.2021.01.035

van Es, M. W. J., Gohil, C., Quinn, A. J., & Woolrich, M. W. (2025). osl-ephys: a Python toolbox for the analysis of electrophysiology data [Methods]. Frontiers in Neuroscience, Volume 19 - 2025. 10.3389/fnins.2025.1522675

Vidaurre, D., Hunt, L. T., Quinn, A. J., Hunt, B. A. E., Brookes, M. J., Nobre, A. C., & Woolrich, M. W. (2018). Spontaneous cortical activity transiently organises into frequency specific phase-coupling networks. Nat Commun, 9(1), 2987. 10.1038/s41467-018-05316-z

Zamanzadeh, M., Verduyn, Y., de Boer, A., Ros, T., Wolfers, T., Dinga, R., Safar Postma, M., Marquand, A. F., van Wingerden, M., & Kia, S. M. (2026). Normative modeling of MEG brain oscillations across the human lifespan. Commun Biol, 9(1). 10.1038/s42003-026-09825-2

Zhang, J., Kucyi, A., Raya, J., Nielsen, A. N., Nomi, J. S., Damoiseaux, J. S., Greene, D. J., Horovitz, S. G., Uddin, L. Q., & Whitfield-Gabrieli, S. (2021). What have we really learned from functional connectivity in clinical populations? Neuroimage, 242, 118466. 10.1016/j.neuroimage.2021.118466

Zhao, Y., Wang, B., Mostofsky, S. H., Caffo, B. S., & Luo, X. (2021). Covariate Assisted Principal regression for covariance matrix outcomes. Biostatistics, 22(3), 629–645. 10.1093/biostatistics/kxz057

